# Polymer Polydispersity and Lipid Composition Control Nanoplastic Disassembly in Membranes

**DOI:** 10.64898/2026.09.24.754055

**Authors:** Hassan Ghermezcheshme, Konark Bisht, Semen Yesylevskyy, Himanshu Khandelia

## Abstract

Nanoplastic interactions with cell membranes can influence particle uptake, translocation, and biological effects, making their molecular description important for assessing the environmental and biological consequences of plastic pollution. However, molecular simulations generally represent nanoplastics using monodisperse polymer chains, whereas nanoplastics formed by environmental degradation are polydisperse. Here, we construct 4 nm polyethylene terephthalate (PET), polyethylene (PE), and polystyrene (PS) nanoparticles with lognormal molecular-weight distributions reported for degraded polymers, and with melt-like chain packing and entanglement. Using Martini coarse-grained molecular dynamics, we simulate their interactions for 10 μs with three lipid bilayers of increasing complexity. PE and PS insert rapidly and remain largely intact, whereas PET remains surface-associated and progressively releases individual chains. Fluid, polyunsaturated lipid environments undergo greater deformation and promote substantially more chain release, up to 56% of the chains for PET and 43% for PS. Release begins with the shortest chains and progressively extends to longer chains, while semicrystalline PE remains largely intact. In contrast, a size-matched monodisperse PS particle releases no chains and minimally perturbs the membrane. These results identify the molecular-weight distribution as a key determinant of nanoplastic–membrane interactions, and indicate that uniform model particles underestimate the release of polymer chains into biological membranes.

## Introduction

Microplastics (MPs) and nanoplastics (NPs) are recognized as a major environmental concern^1^ and may have consequences for human health.^2^ This concern is worsening because plastic production continues to rise rapidly and is expected to reach around 1 billion tons per year by 2050.^3^ Compared to larger plastic particles, NPs smaller than 50 nm in diameter can more readily cross biological barriers such as the blood-brain barrier, are more likely to accumulate in living cells, have a larger surface-area-to-volume ratio and altogether present a higher risk for long-term toxicity.^4, 5^ Humans are directly exposed to MPs and NPs through table salt, drinking water, and air.^6^ The accumulation of MPs or NPs in the human body can trigger cellular-level responses such as increased oxidative stress, activation of immune inflammatory responses, and cellular damage, which may in turn, lead to health-related problems, including impairments in neurological function and metabolic regulation.^7^

Nanoparticles can enter cells through several pathways, such as passive diffusion, direct disruption of the membrane, and both non-specific and receptor-mediated endocytosis.^8^ Since the interaction between nanoparticles and the cell membrane constitutes the initial stage of cellular uptake, understanding nanoparticle– membrane interactions is essential for elucidating their uptake mechanisms and potential biological impact. Polystyrene (PS) nanoparticles have been used as model NPs in most experimental studies. Particles between 20 and 500 nm cross epithelial cell monolayers and model membranes largely by passive translocation, and are internalized by living cells through both passive and active mechanisms.^9–11^ Particle shape also matters: 20 nm PS nanospheres readily permeate artificial phospholipid bilayers, whereas PS nanodiscs bind to and are retained in the membrane.^12^

Molecular dynamics (MD) simulations at both the all-atom (AA) and coarse-grained (CG) resolutions provide a valuable approach for elucidating the molecular details of NP–membrane interactions. The last few years have seen several such studies, and the overall conclusions are rather similar: hydrophobic NPs insert into bilayers and reside in the hydrophobic core, and individual polymer chains disperse inside the bilayer.^13^ Inside the membrane, however, the polymers behave differently: polypropylene and PS chains disperse throughout the bilayer core, whereas polyethylene (PE) chains remain associated and deform into lens-shaped aggregates.^14^ Semicrystalline PE nanoparticles remain structurally intact within POPC and DPPC bilayers, whereas amorphous PE nanoparticles disperse into the hydrophobic bilayer core.^15^ The perturbation of bilayer properties depends on the size of the NPs and the length of the simulations.^16–18^

These conclusions, however, rest on an oversimplified construction of the NPs. To the best of our knowledge, all previous simulations of polymer NP-bilayer interactions have employed simplified NP models based on monodisperse polymer chain lengths. Real NPs generated through environmental degradation, on the other hand, exhibit broad molecular weight distributions (MWD) composed of both short and long polymer chains. Photo-oxidation and mechanical wear break the polymer backbone over time, so plastic debris collected from the environment is enriched in short chains and oligomers compared with the original material.^19–22^ Such polydispersity can influence the behavior of NPs embedded within membranes, as polymer chains of different lengths are expected to detach differently from the parent NP. For example, one would expect that shorter chains are more likely to detach and diffuse into the bilayer, whereas longer chains may conserve NP integrity. Since both experimental and computational assessments of nanoplastic toxicity rely on model particles, the extent to which such particles represent environmentally degraded nanoplastics remains an open question. This concern is not restricted to simulations. More than 90% of studies on nanomaterial-biological interactions use pristine particles of uniform size and shape, and laboratory-weathered PS particles permeabilize model membranes more than pristine spheres do.^23^ That work concerned particle shape; chain-length distribution is a second property separating environmental from model particles, and its effect has not been examined. Three further limitations compound this. First, the exact protocol used to construct the NP will influence individual chain topology and entanglement, which can influence the structure of the NP and its behavior after insertion into the bilayer.^24^ Second, most existing MD studies of NP-bilayer interactions employ simulation times shorter than 2 μs, ^15, 16, 25^ which may be insufficient to characterize the longer-timescale behavior of NPs embedded within the membrane. Third, the force-field parameters used for these polymers are not always accurate. For example, Martini 2-based CG PE nanoparticles remain fully amorphous after equilibration, whereas the corresponding atomistic models form semicrystalline morphologies, and such artificially amorphous NPs can promote unrealistic interactions with lipid membranes.^15^

Whether a nanoplastic remains intact or releases free polymer chains after contacting a membrane matters for exposure assessment, because released chains are far smaller than the parent particle and may distribute and persist differently within tissues. Here, we address the concerns noted above. We construct PET, PE, and PS nanoparticles with experimentally reported molecular weight distributions, anneal them to a melt-like chain packing, and validate their entanglement against polymer physics. We then simulate their interactions with three lipid bilayers for 10 μs, and compare them with a size-matched monodisperse particle and with chemically modified particles. Our aim is to establish whether the polydispersity of environmental NPs changes the way they interact with membranes, and therefore whether monodisperse models are adequate for assessing their biological effects.

## Materials and Methods

We conduct all MD simulations using GROMACS 2023^26^ and the Martini 2 CG force field.^27^ We adopt the mapping schemes and force-field parameters for PS and PE from the parametrization reported by Rossi and co-workers.^28, 29^ The PE topologies used in previous simulations were generated with Polyply^30^ from the CG PE model of Panizon et al.^28^ Polyply assigns the number of exclusions (nrexcl) as 1, whereas the original model uses nrexcl = 3. We find that this difference has a substantial effect on the resulting NP morphology: nrexcl = 3 produces semicrystalline PE nanoparticles whose structural properties agree much better with atomistic simulations and with the expected behavior of linear PE. We therefore use nrexcl = 3 throughout. For PS, we use the A-type mapping scheme described in their study.^29^ For PET, we develop a new

Martini model, optimized to reproduce the structural properties of PET. The complete parametrization and its validation against AA simulations are described in SI-1. For all lipid molecules, we use the bonded and non-bonded interaction parameters of the Martini 2 CG force field.^27^ We model water using standard Martini water beads, with 10% replaced by antifreeze water beads to prevent artificial freezing.

We construct our NPs carefully, following two principles. First, we adjust the MWD of the NPs to closely resemble that reported for degraded particles in experimental studies.^19–21^ Second, to obtain polymer chain entanglement comparable to that in the polymer melt state, we first equilibrate the system at temperatures above the polymer melting temperature and subsequently cool it gradually to room temperature. This procedure promotes chain interpenetration and, for sufficiently long chains, topological entanglement within the NP, comparable to that of a polymer melt. We will quantify the resulting degree of entanglement in the Results section. We then equilibrate each NP in explicit water, and we consider the system equilibrated once the radius of gyration (*R*_g_) of the NP stabilizes. The complete annealing and equilibration protocol is described in SI-2. We generate charged and oxidized NPs starting from the equilibrated configurations of pristine NPs in water. We modify the bead types of selected sites to represent the chemical changes associated with degradation, and we then re-equilibrate the resulting NPs in water using the same protocol applied to the pristine systems. The specific modification sites, Martini bead types, and molar ratios used for each polymer are described in SI-3.

We simulate two bilayers: a pure POPC (18:1, 16:0) bilayer and a mixed bilayer composed of cholesterol, DPPC (16:0, 16:0), and DAPC (20:4, 20:4) in a 0.2:0.5:0.3 molar ratio, which is known to phase separate into liquid-ordered and liquid-disordered domains.^14, 31^ As described later, we find that the NPs deform the DAPC-rich *L*_d_ regions. To check whether this deformation is driven by the phase separation or by the differences in the properties of the DAPC and POPC Martini models, we also simulate the NPs with a pure DAPC bilayer. We generate the lipid bilayers directly using the Insane tool.^32^ For the NP-bilayer interaction simulations, we position the NP above the equilibrated bilayer with a sufficient initial separation to avoid steric overlap with lipid molecules, solvate the system, and add Na⁺ and Cl⁻ ions to obtain a salt concentration of 0.150 M. We perform production simulations for 10 μs at 310 K and 1 bar. The bilayer dimensions and the remaining simulation parameters are given in SI-5.

## Results and Discussion

### Molecular Weight Distribution

By using lognormal distributions calculated from experimentally reported *M*n and PDI values, our NPs have a range of chain lengths that more closely matches those found in real environmental plastic samples, unlike previous studies, which assume a monodisperse (uniform) polymer chain distribution.^15–17^ In all cases, we generate the MWDs assuming a lognormal distribution of molecular weight (ln M). For degraded PET, we use a number-average molecular weight (Mn) = 5000 g/mol and polydispersity index (PDI) = 2.9.^20^ For PE, we select Mn = 5000 g/mol and PDI = 3.0, corresponding to the lowest reported PDI value for degraded PE.^19^ For PS, we use Mn = 5400 g/mol and PDI = 4.6, reported for environmentally degraded PS under accelerated degradation conditions.^21^

The MWDs of the PET, PE, and PS NPs are designed to approximate experimentally reported distributions of degraded polymers within the size limitations of the simulated systems. All three NPs exhibit broad, right-skewed distributions characteristic of environmentally degraded polymers, containing mixtures of both low- and high-molecular-weight chains (Figure 1a). The extended tails toward higher molecular weights indicate the presence of longer polymer chains that preserve the polydisperse nature of the systems, although their abundance is limited by the finite NP size (∼4 nm radius). The charged or oxidized NPs retain particle integrity after chemical modification. The radius of gyration of every NP, pristine or chemically modified, plateaus during equilibration in water, so all particles are structurally equilibrated before contact with the bilayers (SI-4).

**Figure 1.**
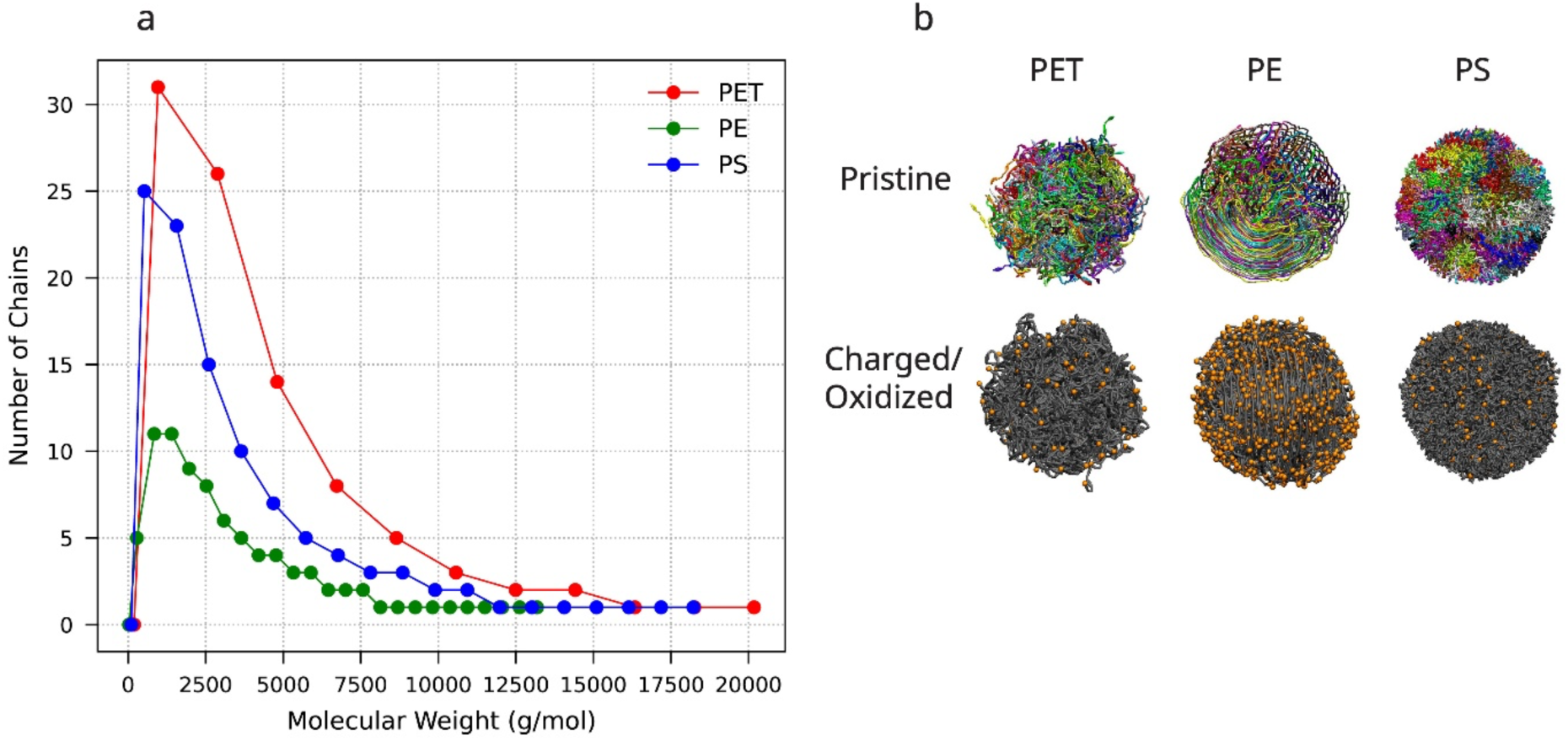
(a) MWD of the polymer chains used to construct the NPs. (b) Representative snapshots of pristine and charged/oxidized NPs. In the pristine NPs, individual polymer chains are shown in distinct colors. Orange beads represent charged or oxidized beads.

### Polymer Chain Entanglement

To characterize the internal topology of the NPs, we perform a shortest-path analysis of the polymer chains using the Z1+ algorithm^33^, as implemented in the open-source code z1plus_neo.^34^ We compute the mean number of interchain entanglements per chain, ⟨Z⟩, averaged over the last 1 μs of the equilibration of each NP in water. We first validate the approach against melt data. For this purpose, we prepare monodisperse NPs of 100-mers using the same annealing protocol, so that they can be compared directly with the Kuhn-scale description of commodity polymer melts^35^, which is formulated for monodisperse chains. In that description the number of entanglements per chain is Z = N_K_ / N_eK_, where N_K_ is the number of Kuhn segments per chain and N_eK_ the number of Kuhn segments between entanglements. Figure 2 compares the Kuhn-scale estimates with our simulated values. Both the magnitude and the ordering of ⟨Z⟩ across the three polymers are reproduced. The agreement is closest for PET and for PE, where our value falls within the reported range. For PS, where the Kuhn-scale estimate is below one entanglement per chain, our value is somewhat higher, although both indicate that 100-mer PS chains are only weakly entangled. Our annealing protocol therefore produces NPs whose internal chain topology is comparable to that of the corresponding polymer melts.

**Figure 2.**
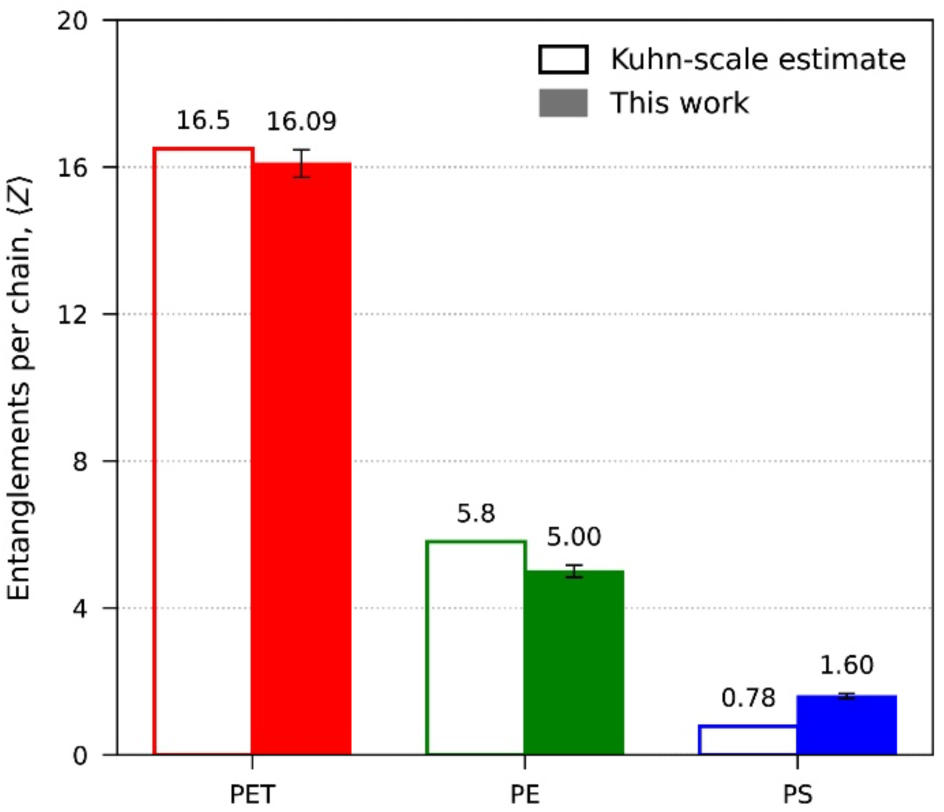
Entanglements per chain, ⟨Z⟩, for monodisperse 100-mer NPs. Open bars: Kuhn-scale melt estimates. Filled bars: this work, with standard deviations.

### Bilayer Equilibration

All bilayer systems are well equilibrated, based on a stable projected area per lipid (SI-6, Figure S-11). For the mixed bilayer, we further confirm the emergence of lateral phase separation into DPPC/cholesterol-rich liquid-ordered (L_o_) and DAPC-rich liquid-disordered (L_d_) domains, which is complete within the first 1 μs and remains stable thereafter (SI-6, Figure S-12). The demixing index used to quantify phase separation is defined in SI-6.^36^

### Interactions of NPs with Lipid Bilayers

Now that we have established the accuracy of our NP models and the equilibration of the membrane, we are ready to describe the interactions of the NPs with the bilayers. Figure 3 presents representative snapshots of the interactions between pristine NPs and lipid bilayers. Movies 1-6 (SI-11) show the temporal evolution of the NP-bilayer interactions. The NP-bilayer interactions depend both on the polymer type and bilayer composition. However, to our surprise, the chemical state of the NPs-pristine or charged/oxidized does not substantially alter the overall interaction behavior.

**Figure 3.**
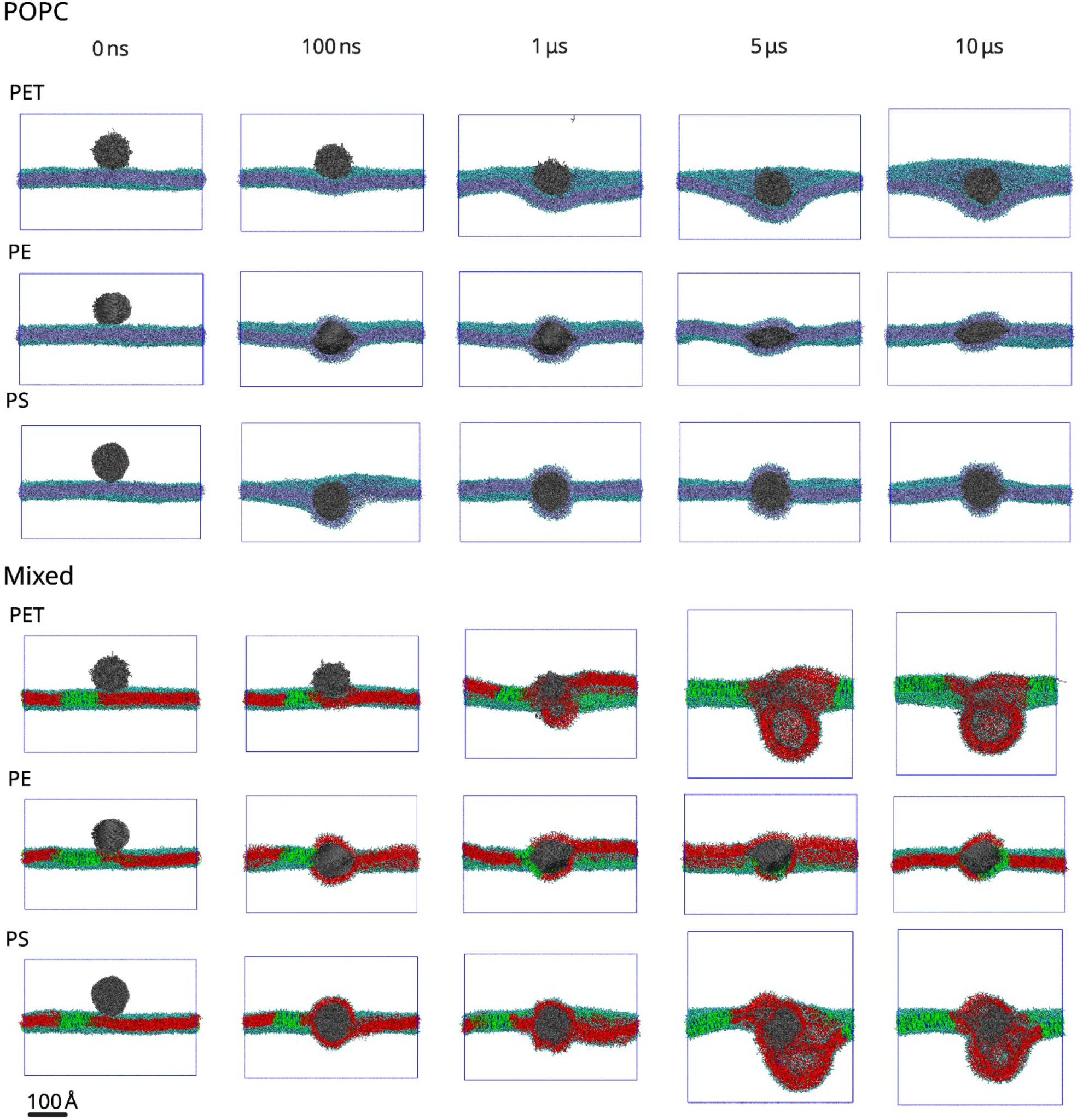
Time evolution of NP-bilayer interactions. Representative snapshots of pristine NPs interacting with pure POPC bilayer and mixed lipid bilayer at selected times. For clarity, the front half of the bilayer is removed to provide a cross-sectional view of the NP-bilayer interface. In the POPC system, lipids are shown in pale blue. In the mixed bilayer, DPPC, cholesterol, and DAPC are shown in green, blue, and red, respectively. Lipid head groups are shown in cyan.

### Interactions with the POPC Bilayer

**PET:** Neither pristine nor charged PET NPs insert into the POPC bilayer within 10 μs. Instead, individual polymer chains gradually detach from the NP surface, with some chains dispersing into the aqueous phase and others embedding into the lipid bilayer. Thus, PET NPs interact with the membrane via progressive chain release and insertion rather than through direct insertion of the intact NP. This behavior is consistent with previous simulations in DPPC bilayers, where PET nanoparticles did not undergo whole-particle insertion into the bilayer.^17^ However, the extended simulation times employed here reveal a progressive chain-release mechanism, in which individual PET chains detach from the nanoparticle and partition into the membrane, a process that was not resolved previously. Individual chain release can locally perturb lipid packing and membrane organization, facilitating the insertion of additional polymer chains. These individual chains may also diffuse laterally within the bilayer and associate with membrane-embedded proteins.

**PE:** Both pristine and oxidized PE NPs insert into the POPC bilayer within 100 ns and remain structurally stable. Upon insertion, the NPs become disk-like (lens-shaped) as seen earlier.^14^ The pristine PE NP deforms more than the oxidized NP, because stronger hydrophobic interactions among the PE chains promote chain alignment and drive a more pronounced morphological transition. Such limited chain detachment was also seen in AA simulations in POPC.^15^

Recently, an improved Martini model of PE was proposed which successfully reproduced the experimentally observed semicrystalline morphology of PE NPs in water.^37^ Once inserted into the membrane, however, the CG nanoplastic spread laterally within the membrane core, dissolved into the lipid environment, and progressively lost its crystalline domains. This is in stark contrast with AA simulations,^15^ where the PE NP became more crystalline, and did not dissolve into the membrane. The PE NP in our CG simulations remains structurally intact. Only a few short chains detach from the NP. Moreover, interactions between the PE chains and the lipid tails promote a more ordered and crystalline NP structure during the simulations, consistent with AA simulations.^15^ Therefore, our PE model reproduces the atomistic behavior of PE NPs interacting with lipid membranes more faithfully than the recently reported Martini 2 model.^37^ The membrane model cannot explain this difference because we use the same Martini 2 lipid parameters. Instead, we attribute the difference to our annealing protocol, which produces melt-like chain entanglement within the NP (⟨Z⟩ = 5.0 for PE, Figure 2), since topological constraints, rather than crystallinity alone, resist the extraction of individual chains.

**PS:** Like PE, both pristine and oxidized PS NPs insert into the POPC bilayer within 100 ns, as expected.^13^ However, unlike earlier studies in which PS NPs lose structural integrity and disperse as individual polymer chains within the membrane core,^13^ PS NPs in our simulations maintain spherical morphology in 10 μs of simulations. This difference arises from the more compact and entangled NP structure generated by our careful preparation protocol. PS is amorphous, so one expects it to retain a spherical morphology.

### Interactions of NPs with phase-separated mixed lipid bilayer

Lipid membranes in cells are laterally heterogeneous and can form functionally-important dynamic domains with distinct lipid compositions, packing, and physical properties. It is therefore important to investigate if NPs preferentially interact with specific membrane environments and how NP can impact membrane phase separation.

#### Visual Inspection

The PET NP approaches and inserts into the mixed bilayer through the DAPC-rich L_d_ domains, where it undergoes progressive chain detachment and partial dissolution, leading to significant local membrane perturbation. The NP inserts into the L_d_ domains irrespective of where they are placed laterally above the bilayer. Previous simulations indicate that PET NPs insert into model mammalian membranes and induce only mild membrane perturbation over 2 μs.^16^

Both PE and PS NPs also enter the bilayer through the disordered domains. The PE NP retains its structural integrity upon bilayer insertion, with no substantial differences observed between the pristine and oxidized systems. Compared to the pure POPC bilayer, PE induces more pronounced bilayer deformation in the mixed system. A similar preference of PE for disordered lipid environments was reported in phase-separated DPPC/DLiPC/cholesterol membranes, where PE aggregates become coated by the L_d_ phase and induce substantial local membrane reorganization.^14^

Since all NPs significantly deformed the DAPC-rich L_d_ phase, we wondered if the deformation was driven by two co-existing phases, or the more fluid properties of DAPC bilayers in the Martini force field. To resolve this, we performed additional simulations of the NPs with a pure DAPC bilayer. The NP-bilayer interactions in the pure DAPC system are qualitatively similar to those in the mixed bilayer, including preferential NP insertion, enhanced membrane deformation, and polymer chain dispersion within the bilayer (Figure S-13). Therefore, phase separation does not lead to increased deformation of the L_d_ phase in the mixed lipid systems. The more pronounced deformation (compared to POPC) results from the higher fluidity of the Martini DAPC lipid bilayer.

### PE Associates more with Cholesterol and Impacts Phase Separation

The three polymers redistribute cholesterol content between the two phases in different ways. We calculate the distribution of cholesterol between the two phases during the simulations, and the impact of the NPs on the degree of lipid phase separation (Figure 4). In every frame, each cholesterol molecule is assigned to the L_o_ phase, the L_d_ phase, or the interior of the NP, based on the lipids surrounding it, as defined in SI-8. Cholesterol is not taken up by any of the particles. The fraction fully enclosed by the NP is zero for PET and PS and below 0.5% for PE, and this category is therefore not shown in Figure 4a. With PE, the cholesterol content of the L_d_ phase increases from 2.3% to 3.6% over 10 µs, with a corresponding decrease in the L_o_ phase from 97.7% to 95.9%. PS does the opposite, its L_d_ content falling to about 1.7%, while PET changes only slightly. PE therefore recruits cholesterol out of the ordered phase, although the effect is modest in 10 µs. Compared to PET and PS, PE interacts more favorably with cholesterol. Linear, apolar PE chains pack against cholesterol much like saturated lipid tails do, whereas the aromatic rings of PS and the aromatic and ester groups of PET make such packing less favorable. A comparable preference of PE for cholesterol-containing lipid environments was reported in previous simulations of phase-separated membranes.^14^

**Figure 4.**
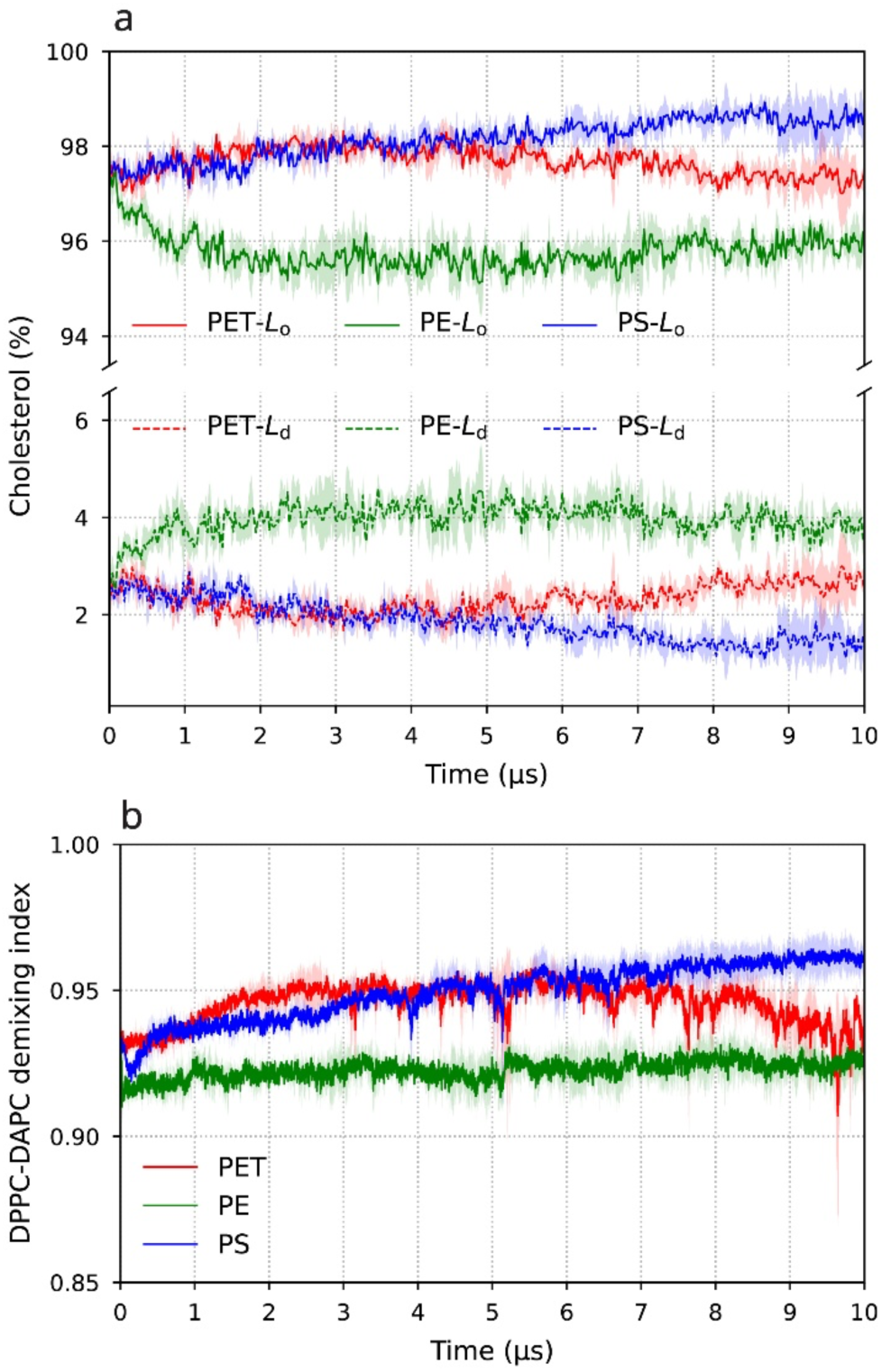
Interaction of PET, PE and PS with cholesterol and their effect on phase separation in the mixed bilayer. (a) Percentage of cholesterol in the liquid-ordered (L_o_, solid lines) and liquid-disordered (L_d_, dashed lines) phases. Cholesterol fully enclosed by the NP is not shown; it is zero for PET and PS and below 0.5% for PE. (b) DPPC–DAPC demixing index during the simulations.

The PE-containing membrane maintains the lowest DPPC-DAPC demixing index, which changes minimally over time (Figure 4b), consistent with the recruitment of cholesterol from the L_o_ phase towards the PE NP in the L_d_ phase. Still, the membrane remains phase separated (*D* > 0.9) in all cases, so the effect on the global L_o_/L_d_ organization is modest in 10 μs. Note that the demixing indices in Figure 4b have not plateaued within 10 μs, so the long-time behavior is difficult to predict from our 10 μs simulations.

### Dynamics of Chain Release into the Bilayer

What is the fate of the polymer chains when a plastic NP binds to the bilayer? If individual polymer chains dissolve into the membrane, they can potentially interact with membrane proteins and influence membrane-mediated biological processes.

To answer this, we calculate the number of polymer chains remaining within each NP (Figure 5), using an algorithm described in SI-9.

**Figure 5.**
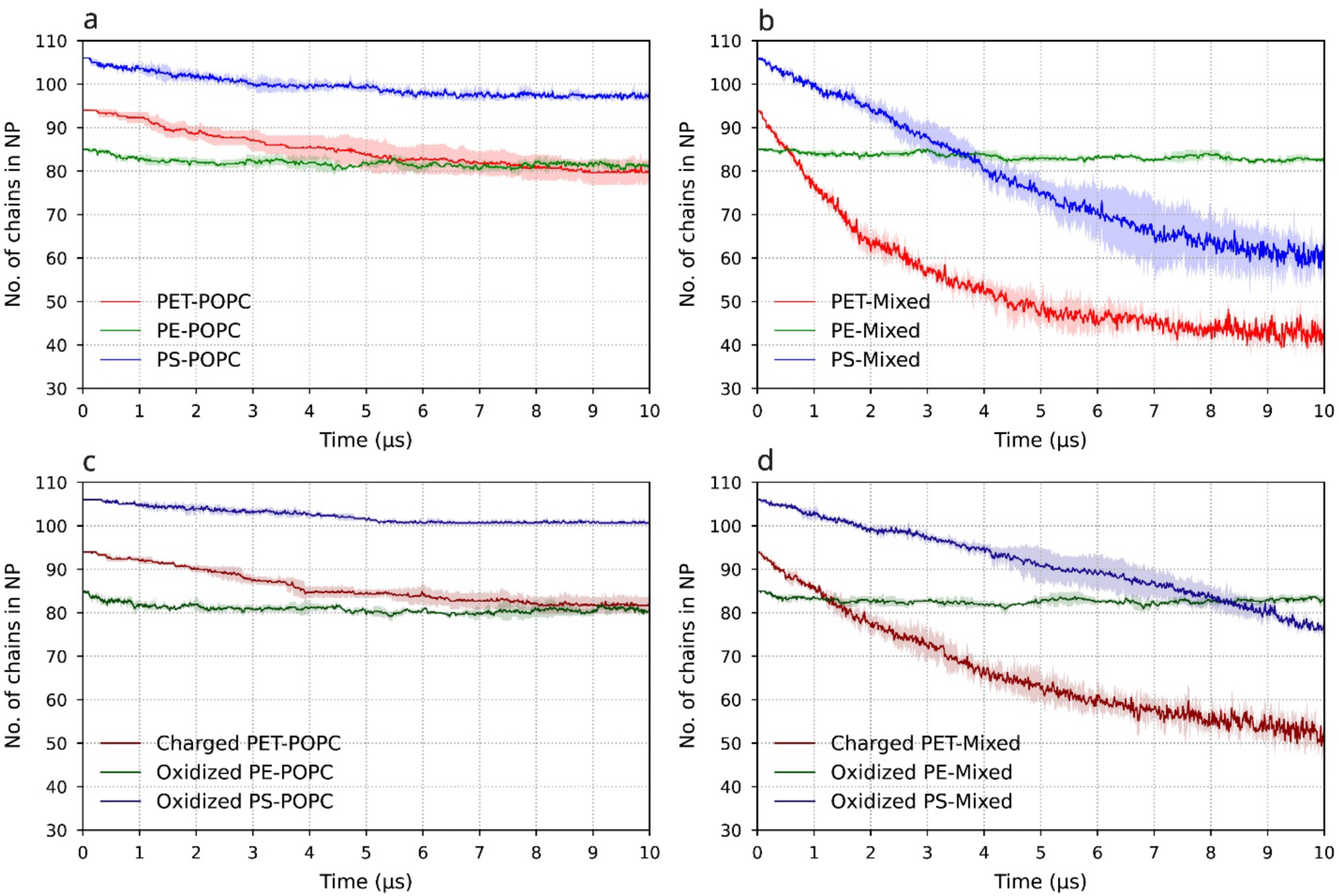
Time evolution of the number of polymer chains remaining in the NPs during interaction with lipid bilayers. Solid lines show the mean of three independent simulations and shaded regions indicate the standard deviation. (a) Pristine NPs in the POPC bilayer, (b) pristine NPs in the mixed bilayer, (c) charged/oxidized NPs in the POPC bilayer, and (d) charged/oxidized NPs in the mixed bilayer.

There is limited chain loss in the pure POPC bilayer. The PE NP loses 4.7 ± 1.0% of its chains and PS loses 7.9 ± 1.6% of the chains. In contrast, PET NPs lose 14.9 ± 3.1% of the chains. PET NPs do not insert into the bilayer, but the polar moieties in PET drive the loss of chains to the aqueous environment and to the lipid bilayer interface. We could not find any differences worth noting between the charged/oxidized NPs and the pristine NPs with respect to NP integrity in the POPC bilayer.

Chain loss is more significant in the mixed lipid bilayer. PS NPs insert through the L_d_ patches. Pristine and charged PET NPs lose 56.4 ± 3.1% and 47.5 ± 2.8% of their chains respectively, and pristine and oxidized PS NPs lose 43.1 ± 2.5% and 26.7 ± 0.9%. The rate of chain release is higher for the pristine PS NP than for the oxidized one (Figure 5b and 5d), which we attribute to stronger hydrophobic interactions of pristine PS chains with the lipid tails. Previously, it was shown that PS chains partition preferentially into the L_d_ phase of phase-separated membranes and promote local redistribution of membrane components.^14^ However, unlike the complete dispersion of PS chains reported previously, the dissolution of PET and PS chains does not reach a steady state within 10 μs in our simulation window. The difference is likely to be a result of our equilibration protocols, which lead to correct chain entanglement, or of the realistic distribution of chain lengths.

The above data triggers three questions. First, why do more polymer chains get dissolved in the mixed bilayer compared to the POPC bilayer? We speculate that the shape of the DAPC-rich environment in the L_d_ phase is altered significantly upon NP insertion, and this can facilitate chain detachment. Second, why are PE NPs more stable within the bilayer? The most obvious explanation is the higher crystallinity of the PE NP, where individual chains pack well against each other and breaking out of this crystalline pattern is energetically unfavorable. Third, what is the chain length distribution of the detached chains? To answer the third question, we analyzed the MWD of the chains released from the parent NP (Figure 6).

**Figure 6.**
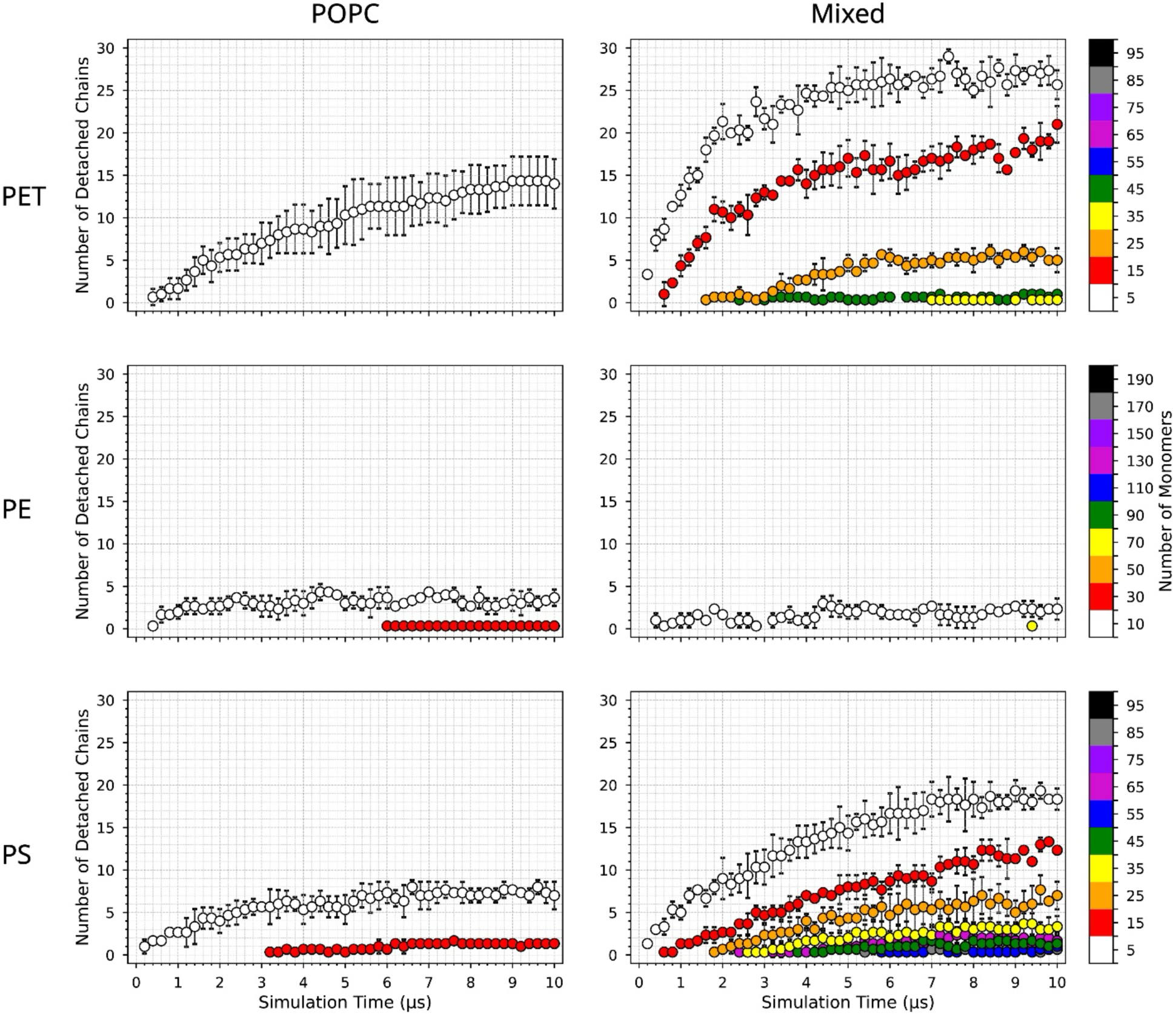
Time evolution of polymer-chain detachment during NP-bilayer interactions.

In the pure POPC bilayer, chain detachment is limited primarily to the shortest polymer chains. For PET, only chains containing 5 repeating units detach from the NP. Similarly, PE NPs release only the shortest chains containing 10 monomers, while PS NPs predominantly release chains containing 5 and 15 monomers. Again, PE releases the fewest small chains due to the semi-crystalline morphology of the PE NPs. In contrast, the mixed bilayer promotes a progressive broadening of the detached-chain population for both PET and PS NPs. Chain release initially involves the shortest chains but subsequently extends to increasingly longer chains as the simulation progresses. Detached PET chains reach lengths of up to 45 monomers, whereas detached PS chains reach lengths of up to 85 monomers. For PE, short chains containing up to 10 monomers get released.

Our data suggest that progressive chain release results in membrane perturbation. In the mixed bilayer, both PET and PS NPs preferentially insert into the bilayer through the L_d_ domains, where the initial release of short polymer chains is followed by the gradual detachment of longer chains, accompanied by progressive disruption of the lipid bilayer. This changes what the NP presents to the membrane. A particle that loses half of its chains delivers free polymer chains into the bilayer, instead of acting as a single object. Weathering is known to fragment plastic not only into smaller particles but also into oligomers and molecular fragments,^38^ which are increasingly recognized as a class of plastic-derived pollutants in their own right.^39^ Our results suggest that these species can also reach a membrane directly from the particle, without first leaching into the surrounding water. To further examine the role of chain detachment, we perform an additional control simulation using a monodisperse PS NP composed of 150-mer chains with the same size as the realistic polydisperse NP. We investigate the interaction of this NP with a pure DAPC bilayer. In contrast to the polydisperse PS NP, no polymer chains detach from the monodisperse NP during the 10 μs simulation, and no significant membrane perturbation is observed. This is direct evidence that the progressive release of short polymer chains, enabled by a realistic MWD, is a key mechanism underlying NP dissolution and the subsequent disruption of the L_d_ phase of the lipid bilayer. Weathered PS particles have been reported to cause greater leakage from giant unilamellar vesicles than pristine spheres of similar size.^23^ There the difference arose from particle shape; here it arises from the chain-length distribution alone, at constant particle size and shape. Uniform particles therefore lack the short chains that start this process. Model particles built from chains of a single length are expected to underestimate how strongly environmentally degraded NPs interact with membranes.

### Location of the Polymer Chains across the Bilayer

Where do the dissociated polymer chains end up in the bilayer? To answer this, we analyzed contacts between the polymer beads and bilayer beads. (Figure 7).

**Figure 7.**
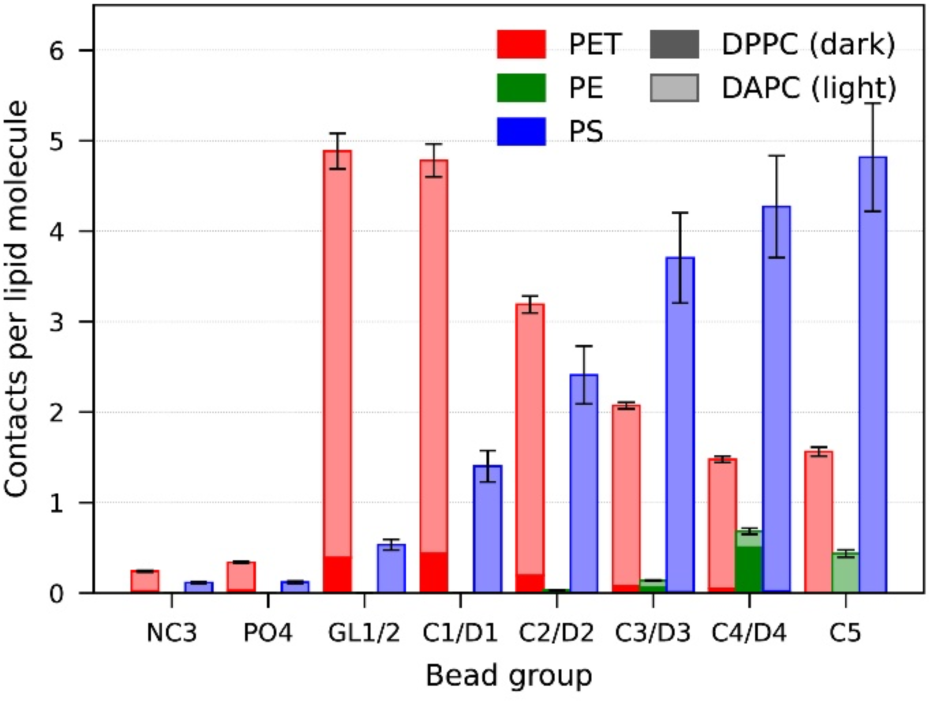
Contacts between polymer beads and the lipid bead groups of the mixed bilayer, normalized per lipid molecule. Dark shading denotes DPPC and light shading DAPC. Bars are the mean over three replicas, each averaged over the last 1 μs, with the standard deviation across replicas. Full definition in SI-10.

The three polymers sit at different depths in the bilayer. PET makes most of its contacts at the glycerol backbone and the first tail bead. On the other hand, the contacts of PS increase with depth and are highest at the terminal tail beads. Thus, the released PS chains partition into the hydrophobic core. PE makes far fewer contacts and only with the deepest beads, as expected for a particle that stays intact and releases very few chains.

Separating the contacts by lipid type shows a second difference. PS interacts almost only with DAPC at all depths, so the PS chains stay within the liquid-disordered phase. PET also makes most contacts with DAPC, but keeps some DPPC contacts at the glycerol backbone and the first tail bead. Thus, when PET reaches the ordered phase, it stays at the interface and does not enter the core. PE has the opposite preference: PE has more contacts with the DPPC lipid tail beads (C4/D4 group) than the DAPC tail beads (C4/D4) although DAPC surrounds the PE NP in the disordered region. The snapshots (Figure 3) show why: the PE particle sits at the boundary between the ordered and disordered domains, with one face in direct contact with the DPPC-rich region. Thus, a large part of the PE NP surface is exposed to saturated tails. This also explains why PE recruits cholesterol from the ordered phase, since cholesterol is concentrated in the ordered domain next to the particle.

Our simulations amount to 0.6 ms of aggregate coarse-grained sampling of three NPs with different lipid bilayers, using equilibration protocols designed to reproduce realistic chain entanglement and experimental chain-length distributions. Taken together, they show that the chemistry of the NP determines both its structure in the bilayer and its impact on the bilayer; that NPs perturb the more fluid, polyunsaturated DAPC bilayer most strongly, and that greater perturbation accompanies greater loss of individual chains; that short chains detach preferentially and act as initiators of further release, producing a cascade that a monodisperse particle cannot reproduce; and that PE interacts preferentially with cholesterol while maintaining the lowest DPPC–DAPC demixing index. We see no major differences between pristine and chemically modified NPs.

Our 10 μs trajectories are longer than previous simulations of NP-membrane interactions. Nevertheless, polymer-chain release from PET and PS NPs, cholesterol redistribution, and changes in lipid demixing were still evolving at the end of the simulations, so our results describe the interaction pathways and relative behavior observed within the 10 μs window rather than the final equilibrium state. Within that window, monodisperse models composed exclusively of long chains suppress the chain-release pathways that occur when short chains are present. Molecular-weight distribution and internal particle topology should therefore be treated as explicit model parameters in simulations of nanoplastic-membrane interactions, and model particles built from uniform chains are unlikely to represent the behavior of environmentally degraded nanoplastics.

## Supporting Information

Martini parametrization of polyethylene terephthalate and validation against all-atom simulations; annealing and equilibration protocol for the nanoparticles; chemical modifications of the charged and oxidized nanoparticles; radius of gyration of the nanoparticles in water; bilayer dimensions and simulation parameters; equilibration, area per lipid, and demixing index of the lipid bilayers; interaction of pristine nanoparticles with a pure DAPC bilayer; assignment of cholesterol to the liquid-ordered phase, the liquid-disordered phase, and the nanoparticle interior; definition of the nanoparticle and of detached chains; contacts between the polymer chains and the lipid bead groups; monodisperse polystyrene control nanoparticle (PDF).

Movies of the interactions between pristine nanoparticles and the POPC and mixed lipid bilayers (MP4).

## Data Availability

All input files for the simulations performed here and representative output is available at Zenodo (DOI: <u>10.5281/zenodo.22936013</u>). The Z1+ analysis was performed with the open-source code z1plus_neo, available at https://github.com/yesint/z1plus_neo.

## Acknowledgment

1. H. Gh. is co-funded by the European Union and the SDU Climate Cluster at the University of Southern Denmark under grant agreement No. 101177011. This work is supported by the Novo Nordisk Foundation Grant number NNF23OC0085169 and by a Villum Foundation Experiment Grant. S.Y. was supported by the grant LUAUS25113 from the Ministry of Education, Youth and Sport of the Czech Republic and the Lundbeck Foundation Visiting Professor grant number R537-2026-114. The MD simulations were carried out on the Danish e-Infrastructure Cooperation (DeiC) on the Finnish Supercomputer LUMI, under grant numbers DeiC-SDU-N5-2024060 and DeiC-SDU-N5-2025105, Novo Nordisk Foundation funded ROBUST Resource for Biomolecular simulations #NNF18OC0032608. This work was also supported by a grant/voucher number NNF25OC0104683 for computational time on the Gefion AI supercomputer from the Novo Nordisk Foundation. We thank the Danish Centre for AI Innovation (DCAI) for operational and technical support.

