## Supporting Information for "Polymer Polydispersity and Lipid Composition Control Nanoplastic Disassembly in Membranes"

### 1. Martini parameterization of polyethylene terephthalate (PET)

In this section, we explain the developed Martini model for the PET polymer. Existing CG models for PET that are not based on the Martini approach have previously been reported.[1, 2] In our CG PET model, five beads represent one monomer unit (Figure S-1). Beads B1 and B4 contain carboxyl groups and are therefore represented using polar small-size SP3 beads. In contrast, B2, B3, and B5 are apolar and are represented using small-size SC1 bead types.

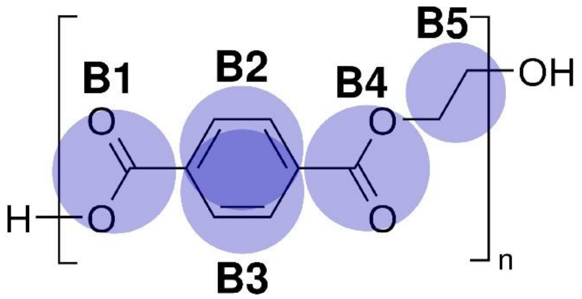


Figure S-1. Martini model for monomer of PET polymer. Beads B1 and B4 are SP3 bead types (small, polar) from the Martini force field, whereas B2, B3 and B5 are represented by SC1 bead types (small, apolar).

We incorporate bonded interaction parameters including bond lengths, distance constraints, and bond angles into the PET model. We define $b_{23}$, $b_{45}$, and $b_{56}$ as the bond lengths between the corresponding beads. To preserve the rigid structure of the benzene ring together with the adjacent carboxyl groups, we apply the distance constraints $c_{12}$, $c_{13}$, $c_{14}$, $c_{24}$, and $c_{34}$. The bond angles included in the model are $a_{145}$, $a_{456}$, and $a_{569}$. A schematic representation of the Martini PET model and its bonded parameters is shown in Figure S-2. We perform extensive simulations to compare the distributions of these bonded parameters in the Martini PET melt with those obtained from an all-atom (AA) PET melt model[3] under identical thermodynamic conditions (see SI-1.1).

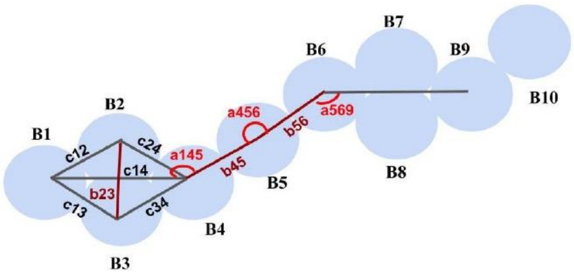


Figure S-2. CG model for PET polymer with different parameters. $b_{23}$, $b_{45}$, $b_{56}$ are bond lengths; $c_{12}$, $c_{24}$, $c_{13}$, $c_{34}$, $c_{14}$ represent distance constraints and $a_{145}$, $a_{456}$ and $a_{569}$ are bond angles between respective beads.

### 1.1. Comparison of macroscopic properties of CG melt with AA melt of PET

PET has a melting point of 523 K and exists in a glassy state at room temperature. Therefore, we perform replica-exchange molecular dynamics (REMD) simulations to sample a broad configurational space for the PET melt.[4] REMD is performed for a melt consisting of 50 PET20 chains using a total of 80 replicas at temperatures ranging from T = 300 K to T = 616 K, with temperature intervals defined according to reference.[5] We compare the macroscopic and microscopic properties of melts described using AA and CG parameters. The simulations are performed for 3 μs, and the averaging is carried out over the last 500 ns. The time evolution of the macroscopic properties is shown in Figure S-3, and the time-averaged values are presented in Table S-1.

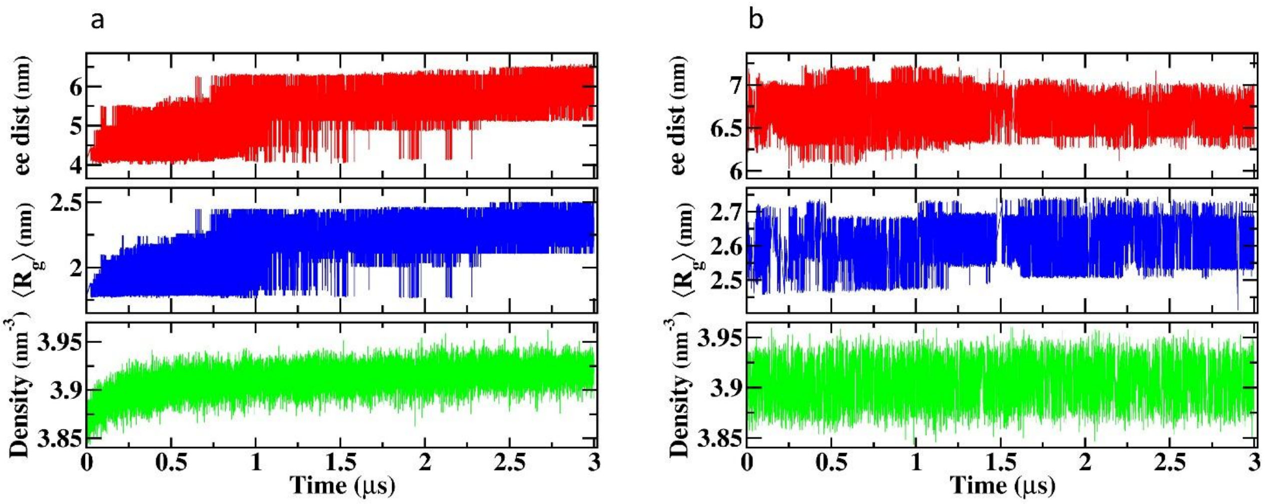


Figure S-3. Time evolution of the end-to-end distance of chains, radius of gyration of chains and number density for (a) AA and (b) CG.

Table S-1. Comparison of macroscopic properties of CG melt with AA melt of PET

| Properties | AA | CG |
|---|---|---|
| Number density (nm$^{-3}$) | $3.919 \pm 0.001$ | $3.906 \pm 0.0001$ |
| Mass density (g/cm$^3$) | $1.251 \pm 0.001$ | $1.463 \pm 0.0001$ [] |
| End-to-end distance (nm) | $5.775 \pm 0.079$ | $6.664 \pm 0.012$ |
| Gyration radius (nm) | $2.297 \pm 0.014$ | $2.600 \pm 0.005$ |

### 1.2. Comparison of microscopic properties of CG melt with AA melt of PET

In this section, we compare the distributions of various model parameters in the CG and AA melts. Figures S-4-6 show the distributions of bond lengths, distance constraints, and bond angles, respectively, for the AA and CG melts.

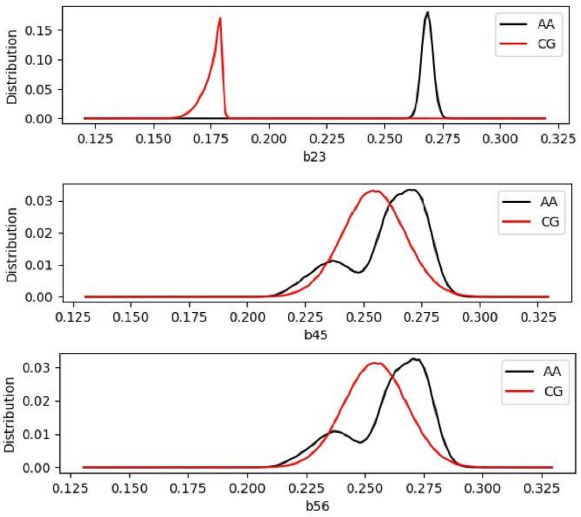


Figure S-4. Distributions of bond lengths $b_{23}$, $b_{45}$ and $b_{56}$ for AA and CG melt.

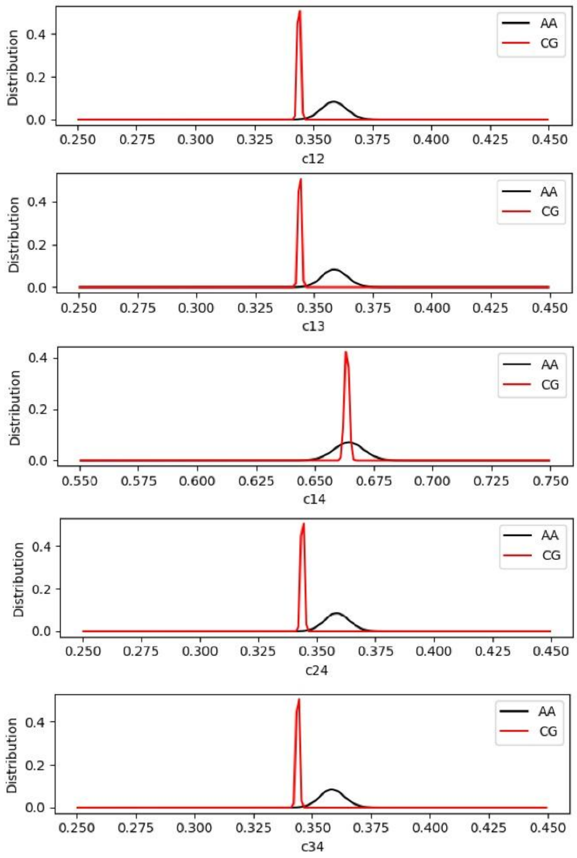


Figure S-5. Distributions of constraints $c_{12}$, $c_{13}$, $c_{14}$, $c_{24}$ and $c_{34}$ for AA and CG melt.

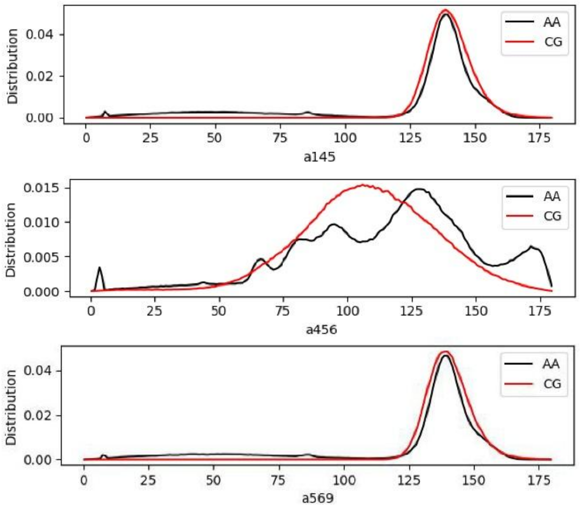


Figure S-6. Distributions of bond angles $a_{145}$, $a_{456}$ and $a_{569}$ for AA and CG melt.

### 1.3. Comparison of macroscopic properties of a single chain of PET in water

We also perform simulations of a single PET chain containing 20 monomer units in water at 300 K. We again compare the macroscopic properties of the AA and CG models (Table S-2). For consistency, the CG simulations are performed using polarizable water (PW).

Table S-2. Comparison of macroscopic properties of single chain of CG PET20 with AA PET20 chain.

| Properties | AA | CG |
|---|---|---|
| End-to-end distance (nm) | $2.149 \pm 0.116$ | $2.171 \pm 0.246$ |
| Gyration radius (nm) | $1.084 \pm 0.032$ | $1.057 \pm 0.0028$ |

### 1.4. Comparison of microscopic properties of a single chain of PET in water

We also compare the microscopic parameters of the PET model for a single polymer chain in water under AA and CG conditions. The corresponding distributions are shown in Figures S7–9.

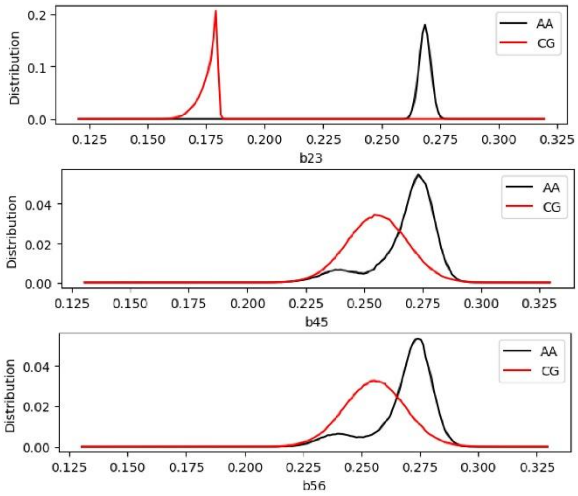


Figure S-7. Distributions of bond lengths $b_{23}$, $b_{45}$ and $b_{56}$ for AA and CG single chain in water.

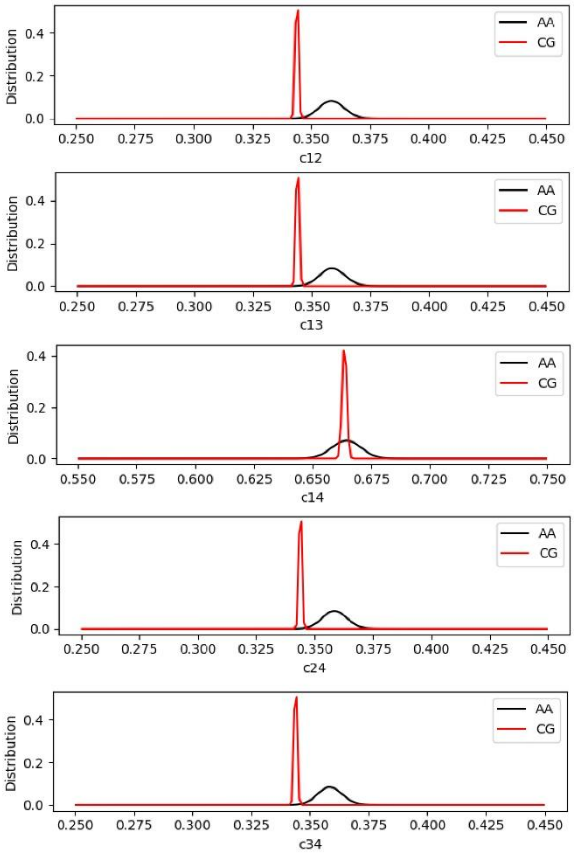


Figure S-8. Distributions of constraints $c_{12}$, $c_{13}$, $c_{14}$, $c_{24}$ and $c_{34}$ for AA and CG single chain in water.

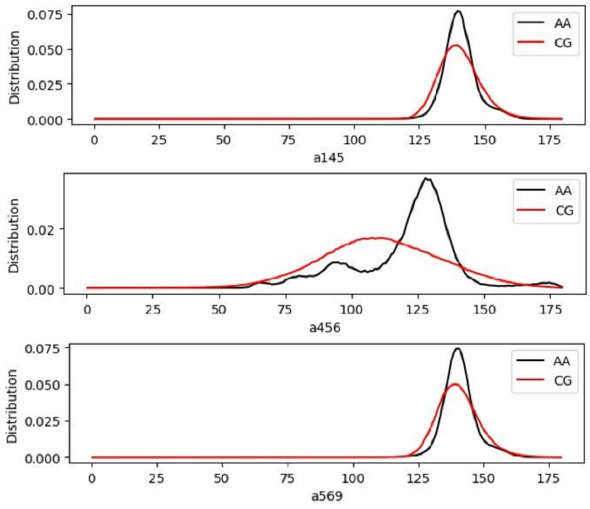


Figure S-9. Distributions of bond angles $a_{145}$, $a_{456}$ and $a_{569}$ for AA and CG single chain in water.

### 2. Annealing and equilibration protocol for the nanoparticles

To obtain polymer chain entanglement comparable to that in the polymer melt state, we first equilibrate the system at temperatures above the polymer melting temperature and subsequently cool it gradually to room temperature. We cool the system stepwise from 2000 K and 0.1 bar to 1000 K and 1 bar in the NPT ensemble, while simultaneously decreasing the temperature and increasing the pressure according to the following protocol: 2000 K-0.1 bar, 1800 K-0.2 bar, 1600 K-0.4 bar, 1400 K-0.6 bar, 1200 K-0.8 bar, and 1000 K-1.0 bar, with 100 ns at each step. Pressure is controlled with the C-rescale barostat, with isotropic coupling, $\tau_p$ = 5.0 ps, and a compressibility of $3 \times 10^{-4}$ $bar^{-1}$. This gradual cooling and compression protocol facilitates progressive intermolecular chain entanglement while reducing premature intramolecular self-entanglement of individual polymer chains before interactions with neighboring chains are established.

We then cool the system from 1000 K to 300 K in the NVT ensemble, in 20 K intervals, and maintain each temperature for 10 ns. In this stage, we reduce the temperature in 20 K intervals and maintain each temperature for 10 ns. After reaching 300 K, we perform an additional 200 ns equilibration without water at 300 K, followed by equilibration in explicit water at the same temperature. We consider the system equilibrated once the radius of gyration ($R_g$) of the NP stabilizes. We use a time step of 2 fs from 2000 K to 300 K, to ensure numerical stability at elevated temperatures, 10 fs for the equilibration in vacuum at 300 K, and 20 fs for the equilibration in water.

**3. Chemical modifications of the charged and oxidized nanoparticles**

For the charged PET NP, we modify the terminal carboxylic acid bead located at the beginning of each polymer chain by assigning a Qa-type Martini bead carrying a −1 charge.

For the oxidized PE NP, we assume that oxidation converts a fraction of C–C bonds into carbonyl groups. We introduce one carbonyl group for every 12 monomer units, distributed randomly along the polymer chains. We randomly select these oxidized sites and modify their bead type to the Na-type Martini bead, representing the presence of carbonyl functional groups.

For the PS NP, we obtain the chemical moieties and their relative abundances from the same experimental study that reports the MWD of degraded PS NPs.[6] We assume that photooxidation produces three main chemical species: carboxylic acids, hydroperoxides, and ketones. We assign molar ratios of 7.5, 6, and 4 for carboxylic acids, hydroperoxides, and ketones, respectively, corresponding to the experimentally measured distribution of oxidation products in weathered PS. These chemical groups are represented using the Martini bead types P2, P3, and Na, respectively.

**4. Radius of gyration of NPs in water**

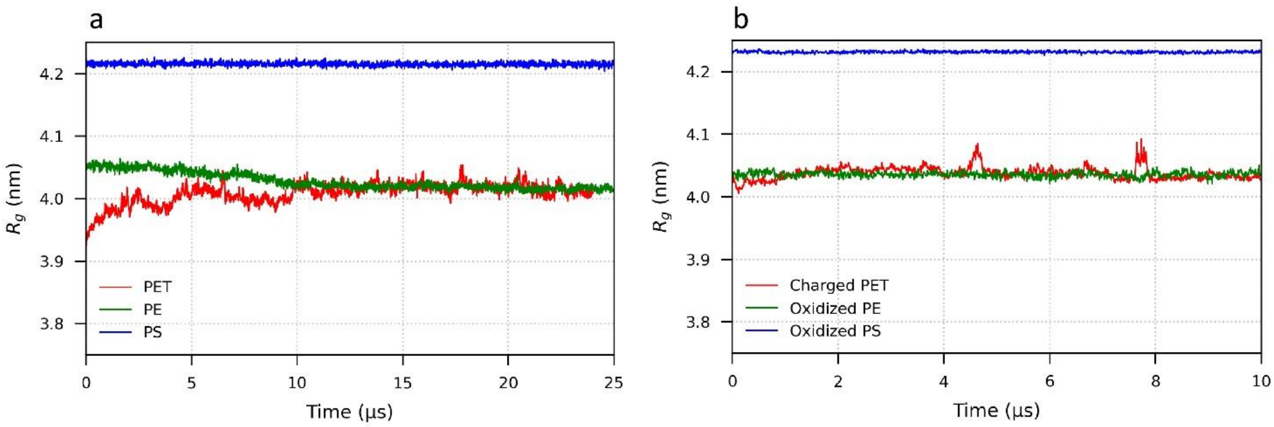


Figure S-10. $R_g$ as a function of time for NPs in aqueous solution: (a) pristine and (b) charged/oxidized NPs.

We evaluate particle stability in water by monitoring the radius of gyration ($R_g$) of the NPs over time (Figure S-10). For PET, $R_g$ increases gradually and stabilizes after approximately 25 μs, indicating moderate swelling and structural rearrangement of the PET NP in water. $R_g$ increases because the polar ester groups in the PET chains interact more favorably with water compared with the fully hydrophobic PE and PS polymers. The $R_g$ of PE decreases slightly before stabilizing, forming a more compact NP structure during equilibration because PE is strongly hydrophobic. The PS NP maintains an almost constant $R_g$ throughout the simulation, consistent with the hydrophobic and amorphous nature of PS Overall, the $R_g$ for all pristine NPs is stable.

For the charged and oxidized NPs, we apply the chemical modifications after equilibration of the pristine NPs in water, followed by an additional equilibration step under the same conditions. The $R_g$ of the modified NPs remains largely stable throughout the 10 μs simulations (Figure S-10b). Thus, the introduced chemical modifications do not destabilize the NPs in water. The small transient increases in $R_g$ at ~5 μs and ~8 μs for charged PET result from the temporary detachment of a few polymer chains from the NP surface into water. Together, these observations indicate that both pristine and chemically modified NPs are structurally equilibrated in water and are ready for the subsequent NP-bilayer interaction simulations.

**5. Bilayer dimensions and simulation parameters**

Each bilayer contains 4224 lipids per leaflet, resulting in lateral dimensions of approximately 51 × 51 nm for the POPC bilayer, 48 × 48 nm for the mixed bilayer, and 59 × 59 nm for the DAPC bilayer. We equilibrate each system at 310 K in the NPT ensemble using the C-rescale barostat ($\tau_p$ = 5 ps) for 5 μs. We apply semi-isotropic pressure coupling with a target pressure of 1 bar, coupled isotropically within the bilayer plane (x–y) and independently along the bilayer normal direction (z).

For the NP-bilayer interaction simulations, we solvate the system using a solvent mixture composed of 90% standard Martini water beads and 10% antifreeze water beads, and add $Na^+$ and $Cl^-$ ions to obtain a salt concentration of 0.150 M. We perform production simulations for 10 μs under the same NPT conditions used for bilayer equilibration.

Each NP-bilayer system is first energy-minimized with the steepest-descent algorithm until the maximum force is below 1000 kJ mol$^{-1}$ nm$^{-1}$. The system is then equilibrated for 1 ns in the NPT ensemble with a time step of 10 fs, and simulated for 10 μs with a time step of 20 fs. Each system is simulated in three independent replicas. All replicas start from the same minimized structure, but with different initial velocities: at the start of both the NPT equilibration and the production run, velocities are generated from a Maxwell–Boltzmann distribution at 310 K with the random seeds 1001, 1002, and 1003 for replicas 1, 2, and 3, respectively.

The temperature is kept at 310 K with the velocity-rescaling thermostat ($\tau_T$ = 1.0 ps),[7] with the NP, the membrane, and the solvent with ions coupled separately. The pressure is kept at 1 bar with the C-rescale barostat,[8] with $\tau_p$ = 5.0 ps. All simulations are performed with GROMACS 2023. The complete protocol, from NP construction to production, is summarized in Table S-3, and all input files are available at Zenodo.

Table S-3. Simulation protocol, from nanoparticle construction to production.

| Stage | Ensemble | T (K) | P (bar) | Barostat | $\Delta t$ (fs) | Length |
|---|---|---|---|---|---|---|
| NP annealing | NPT, isotropic | 2000 → 1000 | 0.1 → 1.0 | C-rescale, $\tau_p$ = 5 ps | 2 | 100 ns per step |
| NP cooling, 20 K steps | NVT | 1000 → 300 | – | – | 2 | 10 ns per step |
| NP in vacuum | NVT | 300 | – | – | 10 | 200 ns |
| NP in water | NPT, isotropic | 300 | 1 | Parrinello–Rahman, $\tau_p$ = 3 ps | 20 | until $R_g$ is stable (25 µs) |
| Bilayer equilibration | NPT, semi-isotropic | 310 | 1 | C-rescale, $\tau_p$ = 5 ps | 20 | 5 µs |
| NP-bilayer minimization | – | – | – | – | – | steepest descent |
| NP-bilayer equilibration | NPT, semi-isotropic | 310 | 1 | C-rescale, $\tau_p$ = 5 ps | 10 | 1 ns |
| NP-bilayer production | NPT, semi-isotropic | 310 | 1 | C-rescale, $\tau_p$ = 5 ps | 20 | 10 µs, 3 replicas |

### 6. Equilibration and phase separation of the lipid bilayers

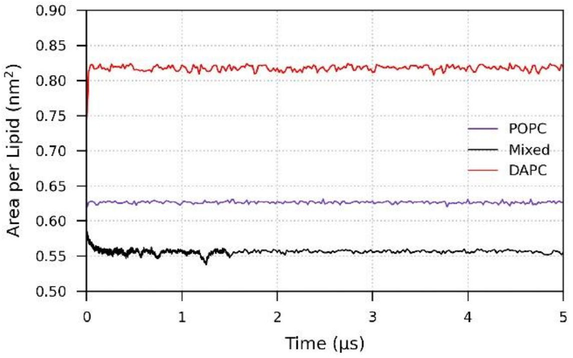


Figure S-11. Time evolution of the APL for the pure POPC bilayer, mixed lipid bilayer and pure DAPC bilayer.

All bilayer systems are well equilibrated based on a stable projected area per lipid (APL) (Figure S-11). For the mixed bilayer we also check for the emergence of lateral phase separation.

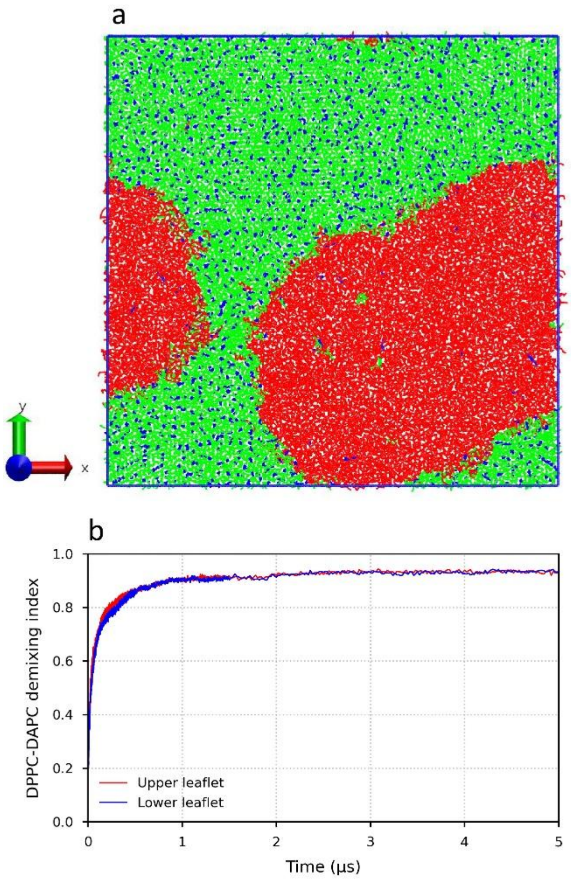

Figure S-12. Phase separation in the mixed lipid bilayer. (a) Snapshot showing the segregation of DPPC/cholesterol ($L_o$) and DAPC ($L_d$) domains; DPPC, cholesterol, and DAPC are shown in green, blue, and red, respectively. (b) DPPC–DAPC demixing index as a function of simulation time for the upper and lower leaflets.

To quantify lateral phase separation, we calculate a normalized contact-based demixing index for each leaflet, following the contact-fraction approach introduced for Martini ternary lipid mixtures.[9] Representative beads (PO4 for phospholipids and ROH for cholesterol) are used to construct a two-dimensional lipid neighbor network using a 1.2 nm cutoff under periodic boundary conditions. We normalize the observed number of DPPC–DAPC contacts by the number expected for a randomly mixed leaflet, so that the index vanishes for random mixing:

$$D = 1 - \frac{N_{\text{obs}}^{\text{DPPC-DAPC}}}{N_{\text{edges}}\, 2 f_{\text{DPPC}} f_{\text{DAPC}}} \qquad (1)$$

where $N_{\text{obs}}^{\text{DPPC-DAPC}}$ is the observed number of neighboring DPPC-DAPC contacts, $N_{\text{edges}}$ is the total number of neighboring lipid pairs, and $f_{\text{DPPC}}$ and $f_{\text{DAPC}}$ are the mole fractions of DPPC and DAPC in the analyzed leaflet. Thus, D = 0 corresponds to a randomly mixed membrane, whereas increasing values of D indicate progressively stronger lateral phase separation. The demixing index rapidly increases during the first 1 µs and subsequently reaches a plateau of approximately 0.93 in both leaflets, confirming the formation of stable DPPC/cholesterol-rich liquid-ordered ($L_o$) and DAPC-rich liquid-disordered ($L_d$) domains.

### 7. Interaction of Pristine NPs with a Pure DAPC Bilayer

Representative snapshots of pristine PET, PE, and PS NPs interacting with a pure DAPC bilayer. All analyses and snapshots of the NP-membrane systems use trajectories processed with the GROMACS tool trjconv: water and ions are removed, molecules are made whole across the periodic boundaries, and the NP is kept as one cluster and centered in the bilayer plane by a translational fit in the *x* and *y* directions. This fit does not change the distances between beads or their positions along the bilayer normal.

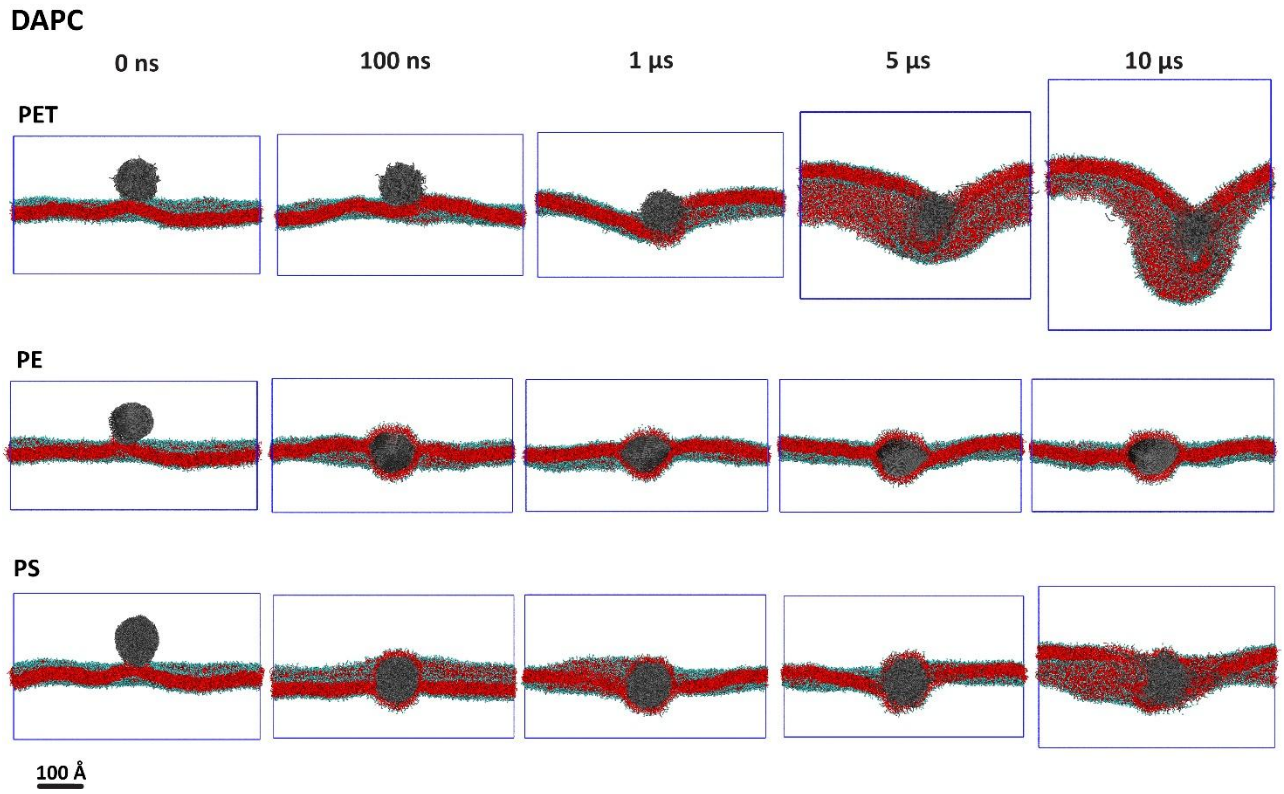


Figure S-13. Time evolution of NP-bilayer interactions. Representative snapshots of pristine NPs interacting with a pure DAPC bilayer at selected simulation times. For clarity, the front half of the bilayer is removed to provide a cross-sectional view of the NP-bilayer interface. DAPC lipids are shown in red, and lipid head groups are shown in cyan.

### 8. Assignment of cholesterol to the $L_o$ phase, the $L_d$ phase, or the nanoparticle interior

In every frame, each cholesterol molecule is assigned to exactly one of three categories. We count the DPPC and DAPC molecules that have at least one bead within 1.2 nm of the cholesterol ROH bead, counting each lipid once regardless of how many of its beads fall inside the cutoff. A cholesterol molecule with more DPPC than DAPC neighbors is assigned to the Lo phase, one with more DAPC than DPPC to the Ld phase, and ties to Lo. A cholesterol molecule with no phospholipid neighbors at all is assigned to the interior of the NP. As a check, we verify that each of these molecules lies within 6 Å of the largest polymer cluster, defined exactly as in the chain-release analysis (Figure 5), so that chains detached from the particle are not treated as part of it.

All beads of each phospholipid are used for the neighbor count, rather than only the phosphate bead. The ROH bead sits roughly 1 nm below the phosphate plane, so a PO4-only network leaves ordinary membrane cholesterol without any neighbors: in the first frame of the PE system, the largest distance from a ROH bead to the nearest PO4 bead is 22.6 Å, whereas the largest distance to the nearest bead of any phospholipid is 5.6 Å. With this definition, a cholesterol molecule resting on the surface of the particle still has phospholipid neighbors and is assigned to the surrounding phase, so that only cholesterol fully enclosed by the particle falls into the third category. No cholesterol is assigned to the NP interior in the first frame of any system, as expected, since the particles are not yet in contact with the membrane.

### 9. Definition of the nanoparticle and of detached chains

For each trajectory frame, we group the polymer beads into clusters using a distance-based connectivity criterion with a cutoff of 6 Å: two beads belong to the same cluster when they lie within this distance, and clusters are built by transitive connection. We define the NP as the largest of these clusters. A polymer chain is considered part of the NP when at least one of its beads belongs to the largest cluster, and is classified as detached when none of its beads is connected to it. The number of chains remaining in the NP therefore provides a direct measure of the structural integrity of the particle and of the extent of chain release.

### 10. Contacts between the polymer chains and the lipid bead groups

We count the contacts between polymer beads and each bead group of the phospholipids in the mixed bilayer. The bead groups follow the Martini lipid structure from outside to inside: the head group (NC3), the phosphate (PO4), the glycerol backbone (GL1/2), and then the tail beads. DPPC has four tail beads (C1 to C4) and DAPC has five (D1 to D4 and C5), so the tail beads are paired by their position along the chain and the C5 group contains only DAPC.

A contact is counted for every polymer bead and lipid bead that are closer than 6 Å. The contacts of each bead group are divided by the number of lipid molecules of that type in the system, 4224 DPPC and 2534 DAPC, so that the values are comparable between the two lipids despite their different abundance. Contacts are counted for all polymer beads, including chains released from the particle. All values are averaged over the last 1 μs of three independent simulations.

### 11. Supplementary movies

Movies 1-6 show the time evolution of the interactions between the pristine NPs and the lipid bilayers over the full 10 μs trajectories. Movies 1-3 show PET, PE and PS with the pure POPC bilayer, and Movies 4-6 show the same three NPs with the mixed DPPC/DAPC/cholesterol bilayer. In all movies the front half of the bilayer is removed to provide a cross-sectional view of the NP-bilayer interface. In the POPC systems, lipids are shown in pale blue. In the mixed bilayer, DPPC, cholesterol and DAPC are shown in green, blue and red, respectively. Lipid head groups are shown in cyan, and the polymer beads in grey. Water and ions are not shown.
